# mcBERT: Patient-Level Single-cell Transcriptomics Data Representation

**DOI:** 10.1101/2024.11.04.621897

**Authors:** Benedikt von Querfurth, Johannes Lohmöller, Jan Pennekamp, Tore Bleckwehl, Rafael Kramann, Klaus Wehrle, Sikander Hayat

## Abstract

Single-cell RNA sequencing (scRNA-seq) transcriptomics improves our understanding of cellular heterogeneity in healthy and pathological states. However, most scRNA-seq analyses remain confined to single cells or distinct cell populations, limiting their clinical applicability. Addressing the need to translate single-cell insights into a patient-level disease understanding, we introduce mcBERT, a new method that leverages scRNA-seq data and a transformer-based model to generate integrative patient representations using a self-supervised learning phase followed by contrastive learning to refine these representations. Our evaluations of mcBERT across 7 million cells from 1223 individuals encompassing diverse disease states in heart, kidney, blood cell, and lung tissues show that learned representations facilitate a robust identification of disease cohorts and enable comparisons of patient similarity. Moreover, our findings indicate that mcBERT can accurately classify disease phenotypes, also in previously unseen biospecimens and patients. Independent of the specific tissue, mcBERT extends the utility of scRNA-seq data from cellular analysis to potentially actionable patient-centric applications.

---

Recent improvements in single-cell technologies have significantly advanced our comprehension of cellular diversity and its implications on human health. This progress is largely attributable to the capability to profile heterogeneous cell populations across diverse modalities at single-cell resolution. Innovative downstream analytical approaches now facilitate detailed examination of gene expression [1], cellular pathways [2], composition [3], and inter-cellular communication [4], fostering not only an enriched understanding of cellular states [5] and disease mechanisms [6–9], but also holding promise for developing therapeutic interventions [10–13].

Furthermore, the application of single-cell methodologies extends beyond cellular and cluster analysis [14]. Indeed, there is a pressing need to develop techniques to draw patient-level conclusions from such analyses, thereby enhancing the clinical utility of single-cell data for patient stratification [15, 16], drug-target discovery [12] and, overall, precision medicine. Although single-cell datasets are inherently high-dimensional, recent advances in data-embedding techniques [17] have succeeded in distilling these complexities into representations that preserve biological integrity on the cell level, such as scGPT [18] and Geneformer [19]. Yet, the potential of such embeddings to also facilitate patient-level comparisons remains underexplored as we lack concrete methods to learn and analyze these representations [20, 21].

Here, we propose a novel method designed to generate patient-level embeddings from single-cell transcriptomics data. Sourcing data from multiple cells and the BERT architecture [22], we refer to this method as multi-cell BERT (mcBERT). Our approach addresses the critical need for a cohesive, patient-centric representation. By employing a novel training pipeline that integrates a transformer encoder and multiple data sources, our method processes individual cell gene counts to produce a compact, disease-capturing patient vector that condenses relevant information learned from single-cell gene expressions. Specifically, the training of our model involves an initial patient-level pre-training phase using a self-supervised data2vec methodology [23], which does not require additional metadata beyond the donor ID. This phase prepares the model to accurately map single-cell data to patient identities, setting the stage for downstream disease-specific comparative analyses. We fine-tune our pipeline using principles from contrastive learning to show that the learned representations are meaningful, specifically, to extract patient-level clinically relevant phenotypical information from individual tissues.

We validated mcBERT across multiple datasets—encompassing over 7 million cells from diverse tissues and pathological conditions (see Table 1)—demonstrates its efficacy in integrating data, mitigating batch effects, and delineating diseases within a disease-oriented latent space. This method to derive meaningful patient-level insights from raw cellular data marks a significant contribution toward the practical application of single-cell technologies in disease diagnosis and treatment stratification.

**Table 1:** All datasets used for the evaluation studies of mcBERT.

| Tissue | Name | #Donors | #Cells | Median<br>#Cells/<br>Donor | Disease Labels |
| --- | --- | --- | --- | --- | --- |
| Heart | litvinukova_2020 [45] | 14 | 199,312 | 13,129 | Healthy |
|  | simonson_2023 [46] | 15 | 89,529 | 6,076 | Long-term Ischemic, Healthy |
|  | kuppe_2022 [6] | 20 | 189,349 | 8,494 | Healthy, Myocardial Infarction |
|  | koenig_2022 [47] | 38 | 216,972 | 5,298 | Healthy, DCM |
|  | chaffin_2022 [48] | 42 | 560,696 | 12,349 | Healthy, HCM, DCM |
|  | reichart_2022 [26] | 70 | 566,809 | 8,007 | Healthy, DCM, ACM |
| Kidney | Muto_2021 [49] | 5 | 19,985 | 3,804 | Normal |
|  | Wilson_2022 [50] | 11 | 39,176 | 2,996 | Normal, Type 2 Diabetes Mellitus |
|  | adpkd1 [51] | 13 | 125,034 | 9,892 | Control, ADPKD |
|  | Kuppe_CD10neg [52] | 15 | 51,849 | 1,493 | Healthy, CKD |
|  | kpmp [53] | 36 | 200,338 | 4,995 | DKD, H_CKD, AKI, Ref, COV_AKI |
| PBMC | PBMC1 [54] | 124 | 836,148 | 6,312 | COVID-19, Normal, Influenza |
|  | PBMC2 [55] | 75 | 422,220 | 5,232 | Normal, COVID-19, Post-COVID-19 Disorder |
|  | PBMC3 [56] | 261 | 1,263,676 | 4,075 | Normal, Systemic Lupus Erythematosus |
| Lung | Human Lung Cell Atlas (Full) [57] | 484 | 2,282,447 | 3,249 | Normal, Pulmonary Fibrosis, Squamous Cell Lung Carcinoma, COVID-19, Lung Adenocarcinoma, Chronic Obstructive Pulmonary Disease, Pulmonary Sarcoidosis, Pneumonia, Lymphangioleiomyomatosis, Interstitial Lung Disease, Cystic Fibrosis, Chronic Rhinitis, Pleomorphic Carcinoma, Lung Large Cell Carcinoma, Hypersensitivity Pneumonitis, Non-Specific Interstitial Pneumonia |
| Total |  | 1223 | 7,063,540 |  |  |

## 1 Results

mcBERT is designed to embed complex transcriptomics data from hundreds of sequenced single cells from an individual patient into a low-dimensional vector that encapsulates the donor’s phenotype. To evaluate our method, we conduct analyses using diverse single-cell datasets derived from multiple tissues (see Table 1), highlighting the broad, universal applicability and flexibility of our approach.

### 1.1 mcBERT Overview

mcBERT starts with the selection of 1023 cells per donor, stratified by cell type, from each tissue dataset (Fig. 1). These cells are characterized by the 1000 most highly variable genes identified across the datasets.

**Fig. 1:**
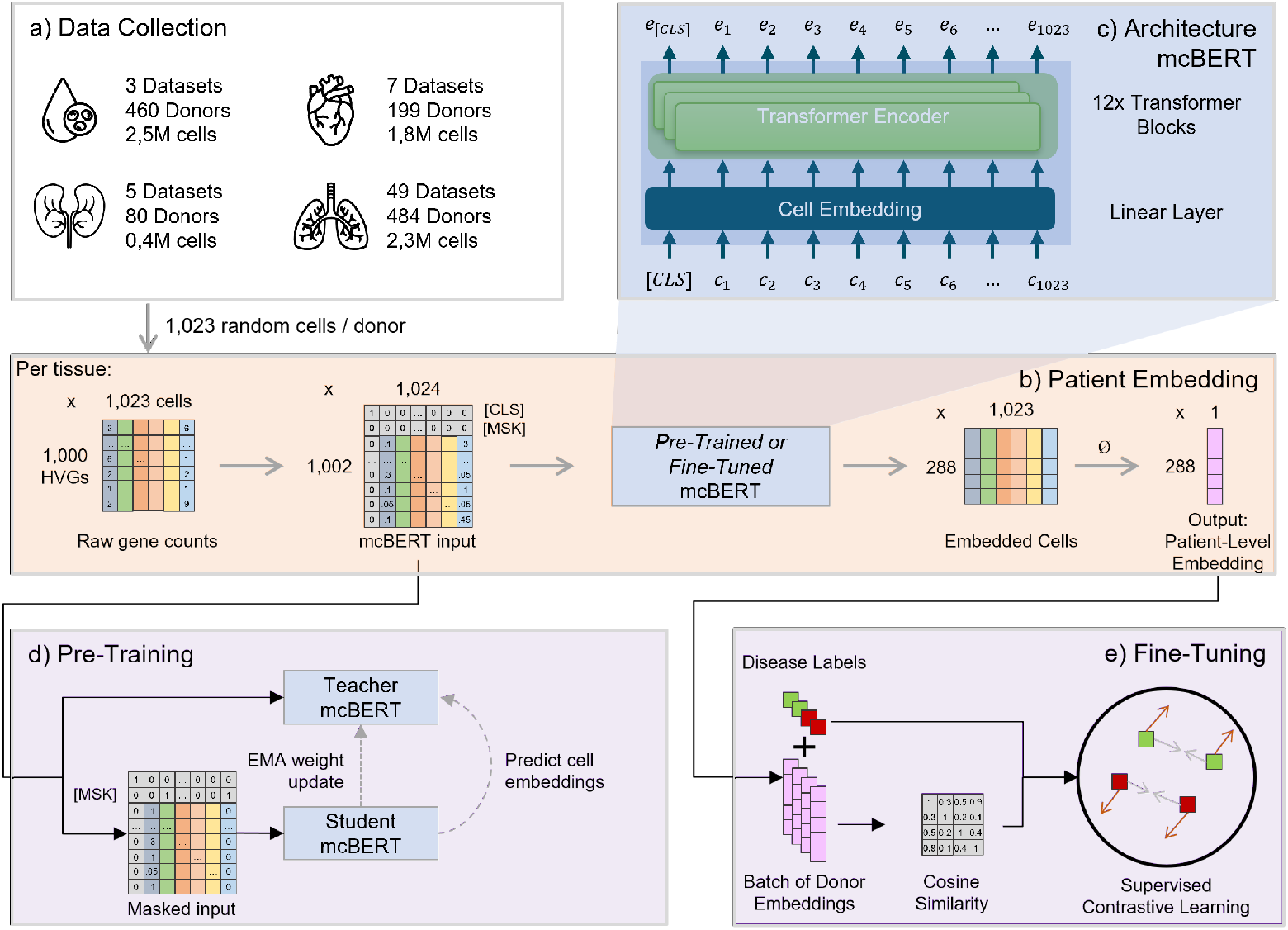
Overview of the patient-level embedding process using mcBERT. a) For training and evaluation, mcBERT utilizes multiple single-cell datasets across four different tissues, for each of which a tissue-specific model is trained. From these, 1023 cells per donor are selected, stratified by cell type, after identifying the most highly variable genes within each tissue-specific dataset. b) The main patient embedding start with 1023 cells represented through the 1000 Highly Variable Genes (HVGs). A classification (CLS) token is added at the beginning, along with special one-hot tokens for masking ([MSK]) and classification purposes. Post-embedding, all cellular data is averaged to generate a singular patient-level vector, which is utilized during inference with the refined model weights. c) mcBERT begins with a non-contextual cell embedding via a linear layer, followed by a transformer encoder composed of 12 blocks, each with 12 attention heads, to contextualize the cell data. d) During pre-training, a student model is fed 1023 cells from a donor, with 15 % of these cells masked. The model’s task is to predict the higher-level representation of these masked cells as determined by a teacher model, which processes the complete, unmasked data. This process is individually conducted for each tissue type, given the variability in HVGs. e) The patient-level embeddings undergo contrastive fine-tuning based on their associated disease labels, employing cosine similarity as the metric for disease similarity. The initial weights for this phase are the tissue-specific weights developed during the pre-training stage.

mcBERT undergoes a two-stage training process on this data. Initially, we employ an unsupervised learning strategy similar to data2vec [23] to capture patient-level correlations across multiple single-cell sequences. Subsequently, the model is fine-tuned using a semi-supervised contrastive learning approach, aiming to cluster donors with similar diseases closer in the embedding space, as indicated by higher cosine similarity.

The efficacy of mcBERT in learning useful embeddings is assessed through its application to various patient-level tasks, illustrating the model’s broad applicability. Primary tasks include evaluating disease similarity directly through the mean cosine similarity across patient embeddings in which, optimally, a cosine similarity of 0 is achieved for patients with dissimilar diseases and a score of 1 with the same disease. Next, precise disease classification based on the local cosine neighborhood of a patient is evaluated via a k-nearest neighbor classifier using accuracy as a score. Finally, the overall clustering quality is analyzed using the Silhouette score [24] to estimate the global clustering qualities ranged from -1 (worst) to 1 (best) based on the separation of the clusters and the Adjusted Random Index (ARI) [25]. Given the number of different diseases of the embedded patients, a hierarchical clustering with average linkage based on the cosine distance first separates the patients into clusters which are then compared with the actual disease labels. The ARI is delimited by 0 for worst clustering results and 1 for best clustering results. These metrics are designed to assess the model’s capability not only to distinguish between diseases but also to generalize across different datasets and diverse biological characteristics of the analyzed tissues.

To validate the resulting model, we begin with experiments on single datasets to establish the baseline effectiveness of patient-level embeddings. Progressively, we increase the complexity of our evaluations by incorporating multiple datasets from varied laboratories to ascertain the robustness and generalizability of mcBERT. This strategy includes testing the model on previously unseen datasets and diseases, thereby verifying its ability to capture and generalize real biological signatures rather than merely memorizing dataset-specific anomalies. Given the utilization of diverse datasets covering multiple tissues, these systematic experiments demonstrate the utility of the learned embeddings for summarizing findings from single-cell data to a patient level.

### 1.2 mcBERT Representation Captures Diseased and Healthy States

We evaluate the capability of raw gene counts from single cells to stratify patients based on disease status using a cardiac tissue dataset comprising 70 donors categorized as Healthy, Dilated Cardiomyopathy (DCM), and Arrhythmogenic Cardiomyopathy (ACM). The dataset is split into training, validation, and testing subsets (70 %, 10 %, and 20 %, respectively) in a stratified way to maintain proportional representation across disease categories. We conduct evaluations under 5-fold cross validation. After training, the validation set is used to determine the best non-over-fitted model and is subsequently used for embedding the donors of the test dataset. For this data, Fig. 2 shows the cosine similarity topology of one fold of the embeddings via UMAP dimensionality reduction.

**Fig. 2:**
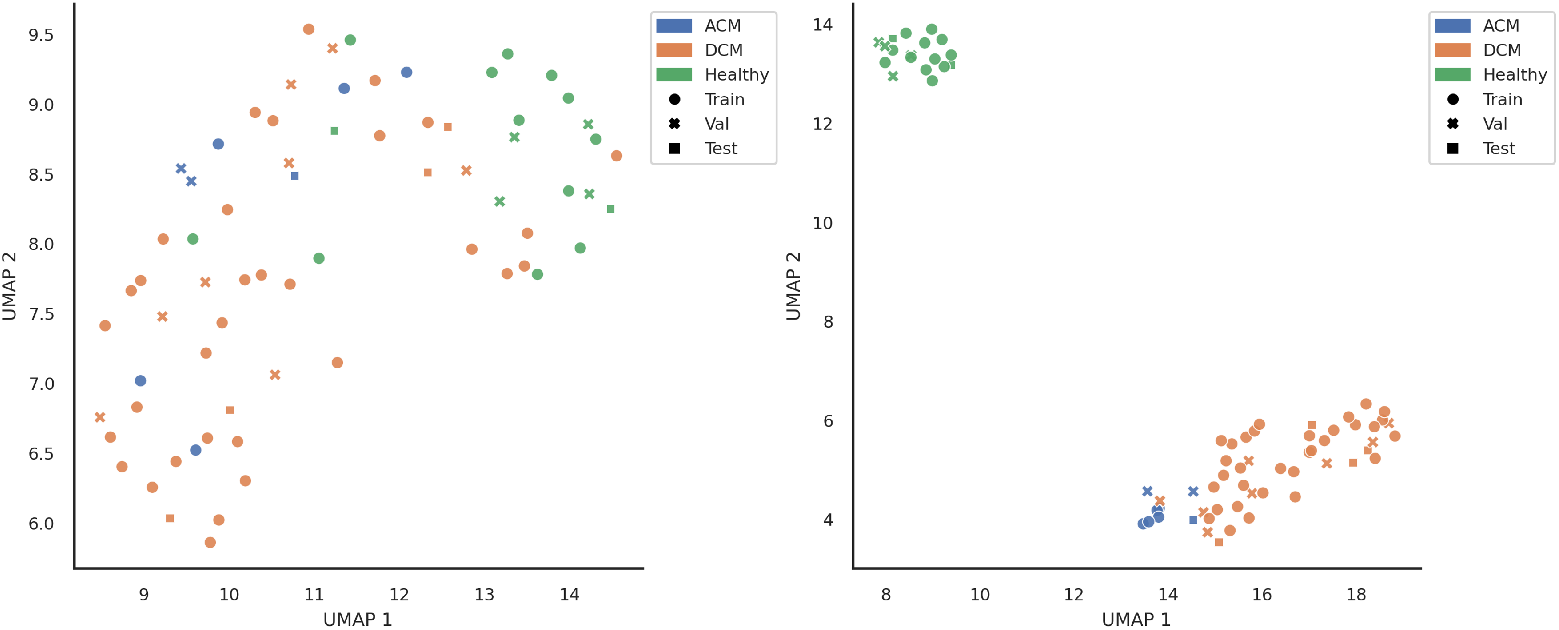
Baseline embeddings (left) versus mcBERT embeddings of the first experiment only using one heart tissue dataset (reichart_2022 [26], 566k cells from 70 donors) visualized using a cosine similarity UMAP. The healthy and diseased patients are well separated, with ACM and DCM forming two sub-clusters among the disease cluster, reflecting the feasibility of patient-centered embeddings using mcBERT.

We compare the performance of mcBERT against a baseline, which utilizes the average cosine similarities of raw gene counts from 1023 randomly selected cells per patient, thereby providing a naïve patient-level representation. The baseline approach exhibits poor clustering performance, with an ARI score of only 0.02 and a Silhouette score around zero, indicating a largely random similarity topology among patients. This is further evidenced by the high cosine similarity among patients with different diseases (0.717) that is close to the cosine similarity of patients of the same diseases (0.774), undermining the potential of raw inputs to effectively distinguish between disease states. Conversely, mcBERT demonstrates superior clustering capabilities, achieving an ARI of 0.766 and significantly improved mean cosine similarities within the same disease (0.889) and significantly reduced similarities across different diseases (0.555). The Silhouette score of 0.635 regarding the whole embedding space underlines these capabilities. These metrics illustrate the enhancement in the model’s ability to embed disease characteristics meaningfully.

This first experiment showed that the raw gene counts of 1023 single-cell data potentially exhibit sufficient information to separate healthy from diseased donors, and the proposed mcBERT architecture with the training method is suitable to extract the relevant correlations inside the gene counts. These results are not limited to the heart tissue, but are also consistent with experiments conducted on other tissues. Therefore, in the following, more complex experiments are conducted to explore the limits and potentials of both mcBERT and the training procedure.

### 1.3 mcBERT representation clusters disease-specific samples across datasets

Mixing multiple datasets for training single-cell models requires addressing inherent challenges such as inconsistent cell-type annotations and technical batch effects [27]. To assess mcBERT’s data integration capabilities, we use the union of all our heart tissue datasets and only standardize cell-type annotations but otherwise, perform no harmonization of gene counts or further batch correction. We resort to the same stratified-splitting configuration and 5-fold cross-validation as before.

Visualizations of the embeddings colored by disease and dataset, respectively, (see Fig. 3) highlight the baseline’s inability to contextualize disease embeddings beyond local dataset characteristics, evidenced by a ARI score of 0.05 which takes into account the global embedding in contrast to a seemingly well-performing model indicated by a k-NN accuracy of 0.72. This inability underscores the necessity of a more sophisticated approach, such as that offered by mcBERT.

**Fig. 3:**
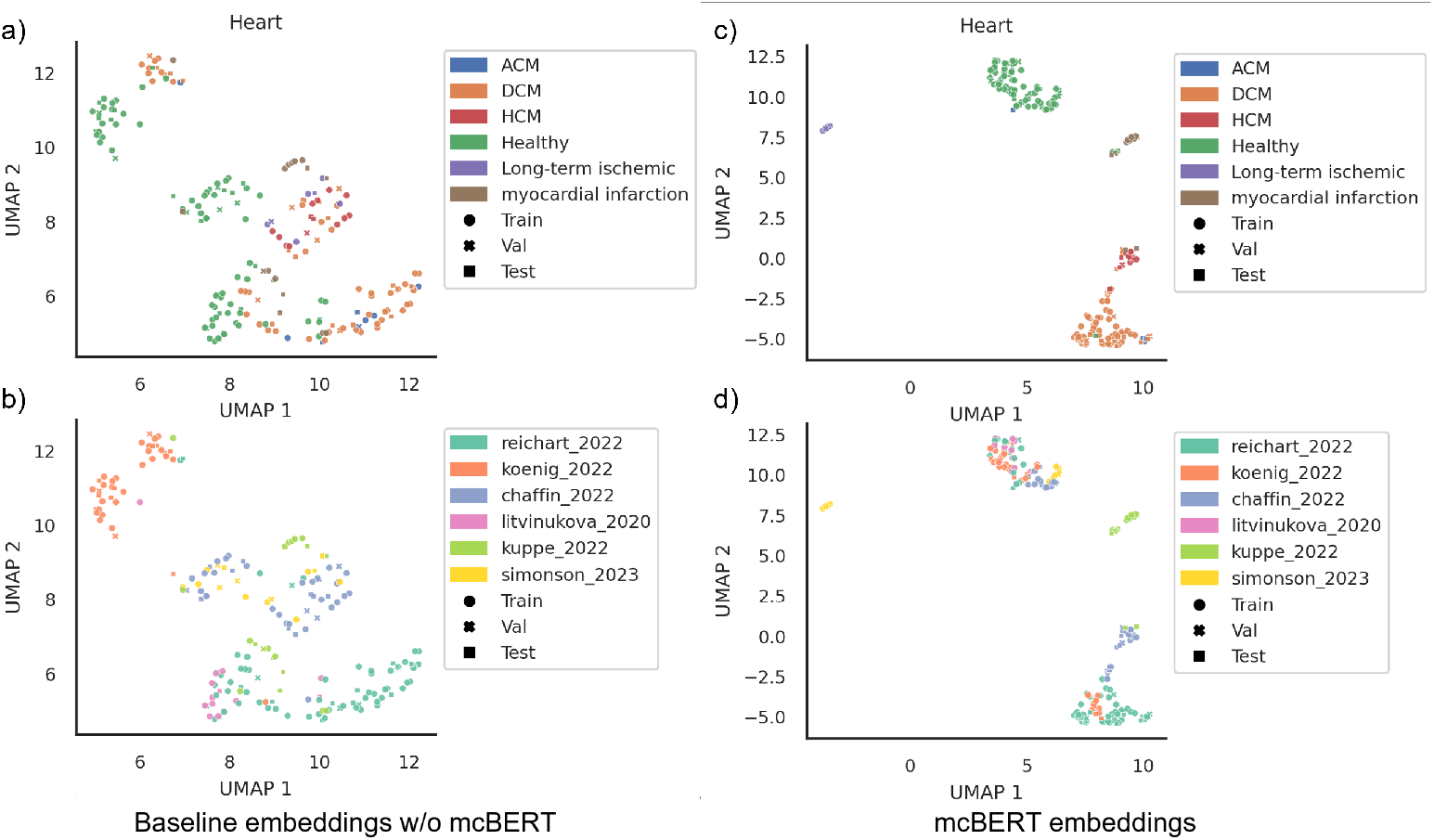
Baseline (a,b) versus mcBERT (c,d) Patient-level embeddings of multiple heart tissue datasets colored by disease (a,c) and dataset origins (b,d). (a,b) shows the baseline embeddings without mcBERT, and (c,d) shows clustering using embeddings obtained by mcBERT. While local dataset-specific clusters are observed in the baseline, mcBERT correctly embeds the diseases across multiple datasets.

Comparatively, the fine-tuned embeddings demonstrate improved clustering across datasets, forming well-clustered groups by disease state, with subclusters for related conditions, such as Hypertrophic Cardiomyopathy (HCM), DCM, and ACM. As these clusters consist of samples from heterogeneous datasets for those disease conditions covered in multiple datasets, this clustering indicates that mcBERT effectively captures and transfers cell-biological knowledge rather than technical batch artifacts or further dataset-specific features.

Applying mcBERT’s pre-training and fine-tuning methodology to samples from kidney, lung, and Peripheral Blood Mononuclear Cell (PBMC) datasets (see Fig. 4) yields consistent results with those from heart tissues, validating our method’s capability to generalize across different biological contexts. Despite the raw input baseline showing relatively high k-NN classification accuracy, significant improvements can be observed in all other metrics. For example, on average, the fine-tuned mcBERT increases the margin of the mean cosine similarity between the same and differently diseased patients from 0.063 to 0.455, demonstrating the model’s effectiveness in overcoming batch effects and integrating multi-tissue datasets. This effect, however, cannot be observed when using the model directly after the pre-training stage, showing both the necessity of the supervised learning stage and the inability to directly derive disease-related information in a self-supervised setting with data2vec for a thorough evaluation of pre-training).

**Fig. 4:**
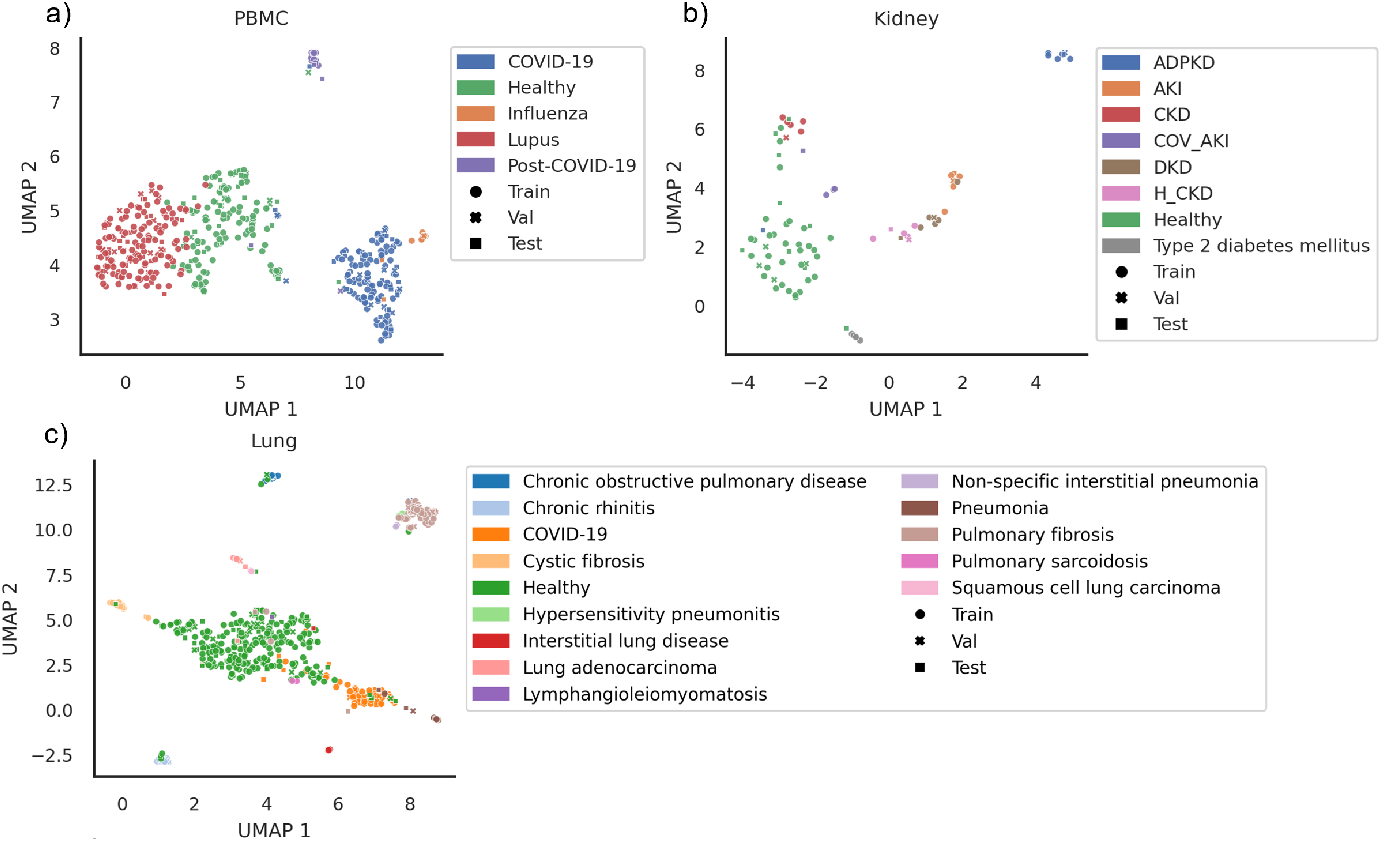
Patient-level mcBERT embeddings of different tissue datasets. a) Kidney, b) PBMC, and c) Lung colored with respect to the patient phenotype. Overall, we observe that mcBERT embeddings can be used to cluster phenotypically similar patients.

To further ascertain the biological relevance of the features learned by mcBERT, we conduct a leave-one-dataset-out cross-validation using six different heart datasets. This approach allows us to assess the model’s ability to apply learned biological information to novel datasets while correcting the batch effects of these datasets. We continue training until observing a plateau in the supervised contrastive validation loss, ensuring that the best model is used for final evaluation. Here, we observe a notable increase in the mean cosine similarity for patients with the same disease (from 0.700 to 0.858) and a significant reduction for those with different diseases (from 0.666 to 0.319), indicating successful disease-specific embedding for entirely unseen datasets.

### 1.4 mcBERT Learns a Meaningful Patient-Level Representation

Typical single-cell analysis frameworks [2, 28–30] use unsupervised clustering techniques to identify groups of cells with potentially similar properties such as cell lineage, cell state [31], and pathway activity [32]. These properties reflect the baseline and dysregulated disease states of groups of cells and are thus vital aspects for disease-centric patient-level neural networks. On a single-cell level, these properties are expressed through e.g., slight changes of gene expressions upon which clustering analyses can be based [33, 34]. To enable mcBERT to learn biologically relevant cell representations, it processes hundreds of cells simultaneously and contextualizes them in the transformer Encoder. Furthermore, the variation in cell type distribution across donors, both natural and introduced through training randomization, ensures that the embeddings generated by mcBERT are not merely reflections of statistical cell distributions.

Besides the transformer consuming multiple cells at once, the training is tailored to the concept of clustering cell types based on the similarity of gene-expression values. It involves a pre-training phase where, through cell masking, mcBERT learns typical gene distributions across multiple cells for different phenotypes. This data2vec-like approach necessitates that the transformer discerns relevant information from surrounding cells to accurately predict the masked cells. This method’s impact is evident when analyzing the integration of the gene-wise dataset; for instance, three independent PBMC datasets initially integrate poorly, showing a scaled iLISI score (larger is better, see the methods section for an overview of metrics) of 0.076. However, post pre-training, this score improves to 0.325 without any dataset-origin information of the donors while maintaining a scaled cLISI score of 0.987 (see Fig. 5). The benefit of this pre-trained cell embedding is underlined by the little to no changes of it after fine-tuning (compare Fig. 5 mid versus right). To effectively predict the phenotypes of the patients, the fine-tuned mcBERT does not separate the cells again to e.g., pick-up dataset-specific batch effects, but relies on using the integrated cell embeddings, emphasizing the biological knowledge extracted.

**Fig. 5:**
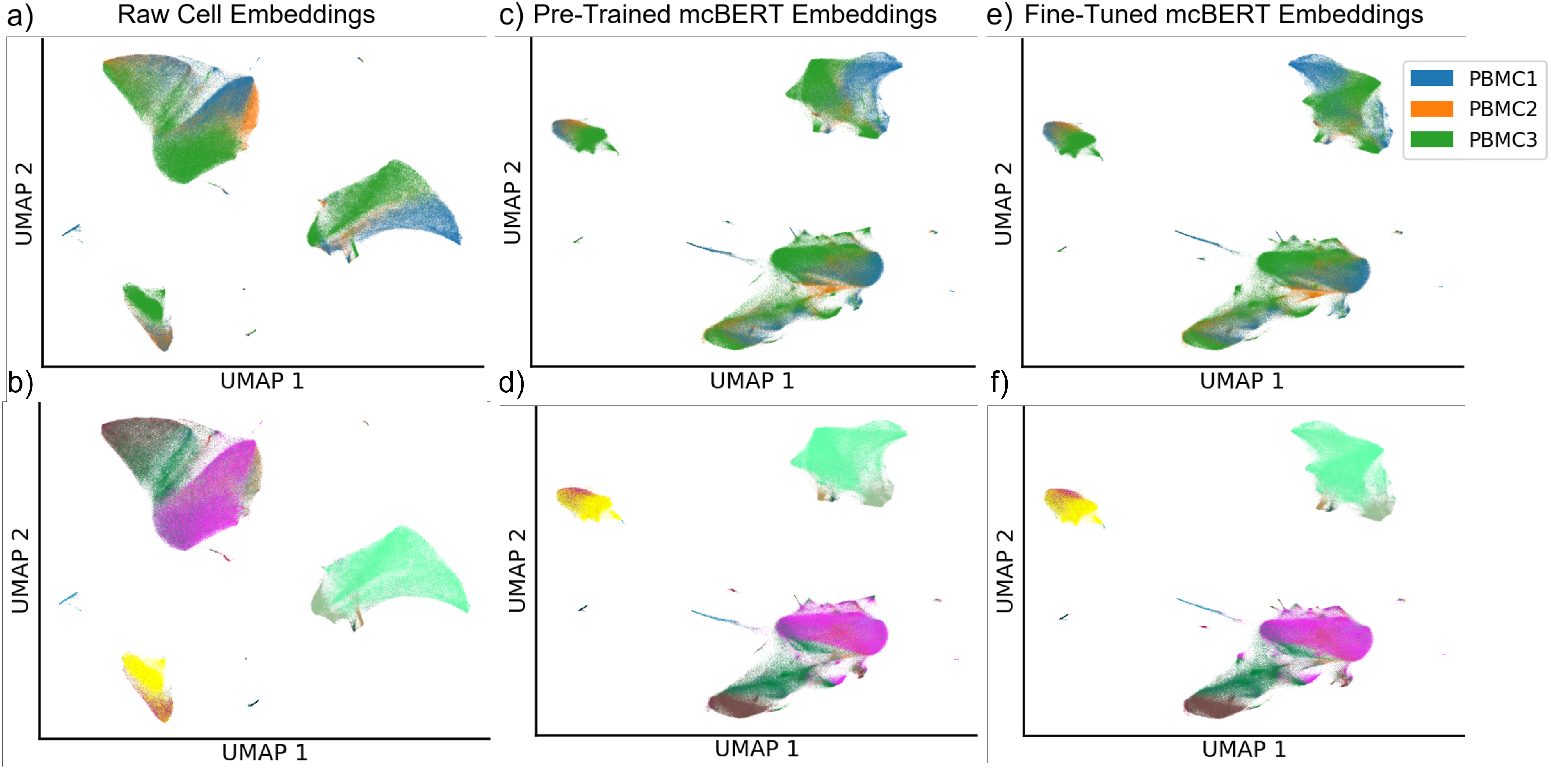
PBMC cell embedding of the first linear layer of mcBERT before training (a, b), after pre-training (c, d), and with subsequent supervised fine-tuning (e, f). Cells are colored by the dataset (a, c, e) and the cell types (b, d, f, legend omitted for clarity). The cell embeddings show clear dataset-wise disparities in the raw gene counts, which are corrected by the pre-training step and are not largely changed with fine-tuning.

Thereby, mcBERT demonstrates its capability to transfer knowledge from samples within one dataset to others at the cell level by embedding similar cell types across datasets in a comparable manner. We attribute this capability primarily to the masking strategy employed during pre-training, which focuses on learning from the cellular context within a donor’s sample to predict the embedding of masked cells, highlighting the importance of this self-supervised training phase.

Additionally, this step further helps preventing early overfitting during fine-tuning and contributes to a more meaningful patient-level embedding. By setting a low learning rate during fine-tuning, the model avoids re-learning dataset-specific technical drifts, ensuring that disease characteristics not present in the training dataset do not skew the embeddings. This approach results in consistent iLISI scores and UMAP visualizations between the pre-trained and fine-tuned phases, as shown in Fig. 5).

### 1.5 Training and Fine-Tuning our Models is Computationally Efficient

We conduct data2vec pre-training for 24 h on a single H100 GPU with 96 GB HBM2e memory, followed by semi-supervised contrastive training with a patience of 20 epochs, monitoring validation loss throughout. In this setting, fine-tuning finishes after approximate 40 epochs without prior data2vec pre-training and around 26 epochs with it. The duration of one fine-tuning epoch varies by dataset, typically ranging from 30 to 130 seconds, depending on the number of unique donors in each tissue dataset. Note that we train tissue-specific models and thus repeat these steps for each tissue, i.e., heart, kidney, lung, and PBMC.

At inference time, mcBERT achieves high throughput, processing over >3100 donors/s using the H100 GPU with 1000 genes and 1023 cells per sample, without further optimization like TensorRT or Flash Attention. Using only a CPU for inference, this throughput drops to 2 donors/s, which we consider sufficient for clinical application. Notably, mcBERT does not require annotated cell types or prior integration of single-cell datasets for inference, enabling efficient edge inference without specialized hardware beyond what is necessary for initial model training and fine-tuning.

### 1.6 Pre-training Boosts Performance of Subsequent Fine-Tuning

Our experiments reveal that mcBERT effectively extracts relevant biological information from individual datasets without requiring their integration. It successfully transfers learned features across different datasets, underscoring its robustness and adaptability. However, the influence of the pre-training and fine-tuning stages on the overall accuracy of the representations generated by mcBERT’s pipeline architecture remains uncertain. To this end, we have already shown that the output of the self-supervised pre-training phase itself is insufficient to carve out phenotypical information of patient level, but it remains unclear whether the fine-tuning alone can already produce accurate representations.

To assess the specific impact of mcBERT’s data2vec-like pre-training, we mirror the approach and chosen methodology but omit the pre-training phase and train mcBERT from scratch directly on the patient-level phenotype separation task. The findings from these tests, as detailed in Table 3, indicate that while tissues from the lung cell atlas show resilience with metrics similar to or slightly degraded from the pre-trained models, the majority yields unstable training behaviors. Specifically, these models struggle to converge, resulting in significantly poorer performance metrics and requiring more epochs to train under identical conditions. These results thus highlight the critical role of data2vec-inspired pre-training in stabilizing the training process of mcBERT and enhancing model performance by effectively harmonizing cell data at the patient level.

### 1.7 Outlook on Clinical Implications

We further show that mcBERT can extract biologically relevant embeddings from hundreds of cells represented by their raw gene counts. This ability generally demonstrates the rich informational content that the sequenced single-cell offers, even with a reduced number of genes of at most the 1000 most Highly Variable Geness (HVGs). If deployed in a clinical environment for, e.g., diseases detection or patient-stratification, focusing only on a minor subset of all possible genes can lower sequencing costs and time, potentially making single-cell practices more accessible and facilitate their broader application.

A clinically relevant tool arising from mcBERT could be a (privacy-preserving) patient-comparison tool: To discover similar disease phenotypes inside a single hospital or a global network of hospitals, one could build upon the patient-level embeddings and integrate these into a search system. Such a tool could offer patient stratification and therapy recommendations for certain phenotypes and give a second data-driven objective view regarding the diseases of a patient without directly relying on sensitive genetic data, thereby improving privacy. While such privacy-preserving properties need further investigation, e.g., using differential privacy on the dimensionality-reduced embedding vector, mcBERT as proposed in this paper, already provides an abstraction layer from sensitive gene expressions.

## 2 Discussion

Traditional scRNA-seq methods typically focus on cell-type level analyses, which leads to challenges in identifying sample-level features and comprehensive sample-vs-sample comparisons. To bypass these limitations, we introduce mcBERT, a novel method that mitigates this situation by abstracting from cell-level to patient-level information via embeddings that represent single-cell data per sample. These embeddings enable patient-level comparisons that can be used to differentiate samples based on their disease phenotype.

Specifically, mcBERT transforms raw, high-dimensional single-cell gene expression data into manageable, low-dimensional representations per patient while preserving biological expressiveness. Thereby, and in combination with advancements from Natural Language Understanding (NLU) and self-supervised learning, mcBERT enables meaningful phenotypical interpretations from single-cell datasets with direct clinical implications.

mcBERT demonstrates robust versatility across tissue types, extracting features from diverse datasets without prior data integration. Notably, the model clusters diseases with similar pathologies closely together, e.g., COVID-19 and flu, and abstracts from batch effects inherent in different datasets without requiring prior integration. We find this methodology to generalize well to novel data and disease classes, offering a significant advancement in the representation of complex, high-dimensional single-cell gene expression data. To our knowledge, mcBERT is the first method to use transformers for systematically deriving patient-level insights from cell-level data. It shows potential for disease and phenotype detection and is readily available (see code). Although the high costs of scRNA-seq limit its routine clinical use, future integration of explainable AI techniques with mcBERT might contribute to an in-depth analysis of the model’s decision-making processes. This analysis helps identify expressive cell types, states, and genes that can then be further utilized as marker genes for accurate and more cost-effective disease testing and drug discovery.

For future work, we plan to expand mcBERT to accommodate multi-tissue single-cell data through a larger foundational model. This expansion is expected to improve the model’s performance and allow for cross-organ insights. Furthermore, leveraging the broad selection of single-cell datasets available, such as cellxgene [35], cancer cell atlas [36] and the human cell atlas [37], will enhance our understanding of biological phenotypes and disease mechanisms, identification of pathogenic cancer cells, proliferation, and early disease detection. Overall, we see immediate application potential of mcBERT and further expect long-term benefits that follow from its patient-focused methodology.

## 3 Online Methods

mcBERT leverages the Bidirectional Encoder Representations from transformers (BERT) framework [22], well-known in Natural Language Processing (NLP). Initially, the model is trained unsupervised to generate concise representations of biological cells and their interrelations, mitigating pronounced batch effects specific to datasets. Subsequently, our training employs contrastive learning to foster a disease-centric comparative analysis of cells from various donors. In the following, we provide a detailed description of mcBERT’s architecture, the processes for training, data preprocessing, and inference, and conclude with details on the employed evaluation metrics.

### 3.1 Approach Design

#### Architecture Overview

Patient-level mcBERT adapts the BERT architecture [22], applying its core NLU principles to the biomedical domain. The model processes multiple input tokens, beginning with a classification token ([CLS], as shown in Fig. 1) (Step 1). Each cell token is then embedded individually in the Cell Embedding layer (Step 2), processed through a transformer Encoder (Step 3), and ultimately pooled to derive a unified patient-level embedding (Step 4). Supplementary Table 2 describes the configuration of these layers.

#### Step 1: Input Selection

Patients are represented by numerous sequenced cells, each characterized by raw gene counts across thousands of genes. Common practice targets a subset, specifically the HVGs, to aid in cell type and state identification [38]. In the initial step, only the *m* most HVGs are selected, reducing input dimensionality and computational complexity. Each cell is represented by an *m*_*genes*_ *×* 1 vector, with each entry reflecting the normalized count of a selected gene. To capture intercellular correlations, *n* cells per patient are randomly chosen, with stratification by cell type to stabilize training and ensure consistent cell type distribution throughout model training and evaluation.

#### Step 2: Cell Embedding

The Cell Embedding layer, modeled after the BERT architecture [22], features a linear transformation with an input size of 2 + *m*_*genes*_ and an output size of 288, simplifying computation in subsequent attention layers. Unlike traditional BERT [22], we omit positional embeddings, making the model positionally invariant. The spatial information of extracted cells is typically not retained during tissue sampling, resulting in few datasets with reliable or standardized positional data for model use. However, if relative cell positioning is crucial and preserved during extraction and sequencing, as in Spatial transcriptomics, incorporating two- or three-dimensional positional encodings into the model remains a feasible adaptation. Related methods like PILOT [20] and Harmony [27] utilize Principal Component Analysis (PCA) for dimensionality reduction [38]. PCA exclusively relies on the statistical properties of the input, the different genes of numerous cells, and represents them through linear combinations. However, our approach replaces PCA with a fully connected layer without activation function, which serves as a learnable dimensionality reduction tool, similar to scBERT [39].

#### Step 3: Transformer Encoder

Until now, all cells have been processed individually. To detect correlations across multiple cells (and thereby genes), they are processed through a transformer encoder. Here, all of the stratified randomly selected cells are processed by multiple self-attention layers, which can extract important correlations across the cells on multiple layers of abstraction.

#### Step 4: Pooling

Post-encoding, *n*_*cells*_+1 embeddings are computed. To consolidate these into a single patient-level vector, we apply global average pooling across all embeddings, a method also employed in vision transformers [40] and offering comparable efficacy to using the [*CLS*] token. The final embedding vector represents the aggregated gene expression profile, prepared for downstream tasks focused on disease-specific embeddings.

#### Number of Genes and Cells per Donor

Selecting an optimal number of genes from each cell and cells from each donor presents significant challenges. To manage computational complexity and memory consumption, we utilize only the 1000 most HVGs, a common practice aimed at reducing overfitting and enhancing neural network speed while potentially sacrificing some donor-specific information. When aggregating single-cell data across multiple datasets, HVGs are identified separately for each dataset, and their intersection is used. To consistently yield 1000 HVGs, we increase the number of extracted HVGs per dataset until their intersection meets this threshold.

The choice of cells per donor to include is driven by the typical cell counts per donor in our datasets, as detailed in Table 1. While more cells per donor might improve the embedding accuracy of the mcBERT network, it slows down both training and inference. Additionally, using random sampling, more cells per donor reduces sample variability, potentially decreasing the diversity of training examples. Given the datasets’ lowest median cell count per donor is 1493, we select 1023 cells per patient to balance performance and computational efficiency.

### 3.2 Model Training

mcBERT undergoes a two-phase training approach. Initially, the model is trained unsupervised to learn the first basic correlations between a patient’s cells and genes. This foundational understanding facilitates the subsequent phase, where disease-oriented embeddings are refined using a supervised contrastive learning strategy.

#### Overview

While the network design allows for the generation of patient-specific vectors, these vectors require initial training to properly embed and correlate cellular information. We thus propose a patient-comparison task where the model learns to differentiate among various diseases and recognize similarities across patients. The objective is for the network to embed patients such that the cosine similarity between patient-level vectors is 1 for patients with the same disease status (i.e., same disease or both healthy) and 0 for those with differing statuses. This approach trains the network not directly on disease identification but rather on recognizing cell and gene combinations indicative of similar or distinct disease states.

Further, training with patient-level comparisons enhances model flexibility in several ways. First, it allows the embedding of disease similarities to guide training, yielding semantically richer and more interconnected embeddings. Second, it enables class-aware semi-supervised learning, where the exact diseases need not be known during training, allowing the model to potentially cluster undiagnosed or novel diseases based on learned patterns. However, prior to this focused training, the model is pre-trained unsupervised to develop a broader understanding of intercellular relationships within individual patients.

#### Self-Supervised Learning

Self-supervised learning generally helps to discover underlying structures and correlations in datasets without a specified training target and is widely used for pre-training single-cell neural networks. Unlike cell-based transformer models that commonly employ masking strategies for training on gene correlations in single cells [18, 39], our approach raises this masking strategy to the patient level, focusing on the training of cell interrelationships in a self-supervised manner. We adopt a masking strategy akin to data2vec [23], aiming for an initial contextual understanding at the patient level involving over a thousand single cells.

The data2vec training regime involves two configurations: a “student mode” and a “teacher mode”. In student mode, 15 % of the input cells are randomly masked with a one-hot encoded masking token ([MSK], see Fig. 1), and the model predicts the representations of these masked cells. The remaining 85 % of cells from the same patient aid in this prediction. The teacher model, in contrast, processes the same inputs without any masking.

Drawing parallels with NLU, the training of the student model utilizes a smooth L1 loss combined with the Adam optimizer and a learning rate of 1 × 10^−5^, following the suggestions of data2vec for text processing [23]. The loss function compares the student’s embeddings of masked cells against the teacher’s embeddings of the same, unmasked cells:

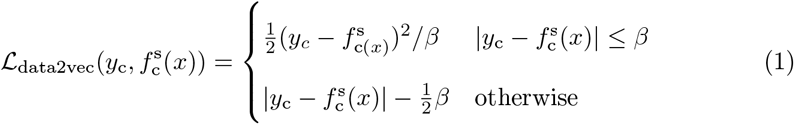

Here, *y*_c_ is the normalized and averaged output of the top 6 transformer encoder blocks for the contextualized embedded cell *c*, and 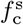 denotes the student model’s output for cell *c*. The threshold *β* for transitioning between L1 and L2 loss in the smooth L1 loss is set to 4, following data2vec NLP recommendations [23].

Subsequently, the teacher model weights are updated using an Exponential Moving Average of the student’s weights, preventing model representation collapse [23]. This self-supervised method, proven effective in other pre-training contexts [41], enables the model to predict masked cell representations accurately using the contextual information from remaining unmasked cells, fostering an understanding of patientspecific cell interrelations. This general understanding allows for further training on more specific tasks.

#### Supervised Contrastive Learning

We conduct patient comparisons by calculating the cosine similarity and distance between embeddings *e*_*A*_ and *e*_*B*_, produced by forward processing the cells from patients A and B. Afterward, the cosine similarity and cosine distance are calculated as follows:

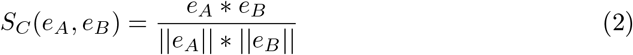

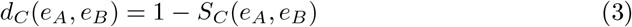

We then apply a contrastive cosine embedding loss function:

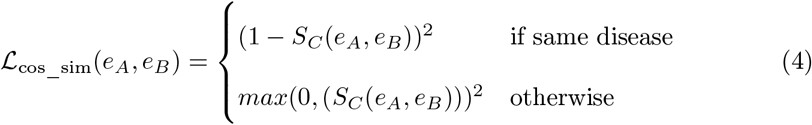

Here, for each patient in the dataset, a second patient is randomly selected with a probability of 50 % that it belongs to the same disease and with another 50 % of a different disease. As stated before, from each patient, *n*_*cells*_ random but stratified against their cell types cells are selected, separately processed by the model, and afterward compared using the loss function.

While this standard contrastive learning setting is commonly applied for, e.g., embeddings of sentences as done in Sentence-BERT [42], it often exhibits unstable training behavior and less global meaningful embeddings, as only two random data samples are compared based on which the network is tuned. When not only the bilateral relationship of the training instances but all general labels (like diseases) are known, one can use all of the labels of one training batch and train on pushing apart patients with different diseases while pulling together the same instances, as proposed with the Supervised Contrastive (SupCon) loss [43]. Given the instability of traditional contrastive learning and the availability of disease labels, we employ the SupCon loss function. Hyperparameter testing indicates that choosing an AdamW optimizer [44], together with a learning rate of 1 × 10^−5^ and a batch size of 48, yields good performance and relatively stable training.

#### Data Sparsity

Today, large open-source single-cell RNA datasets often exceed 1 million sequenced cells, facilitating the development of advanced cell-level deep learning models. However, the number of donors typically falls below 100 per dataset. With a typical test dataset allocation of 25 %, fewer than 75 donors are available for training. Combining multiple datasets from similar tissues and diseases does not sufficiently overcome the challenge of limited donor numbers, exacerbated by varying technical batch effects and a frequent underrepresentation of disease classes compared to a ubiquitous “healthy” reference class. For that reason, large collections of datasets, e.g., in the form of so-called cell atlases, can be especially valuable and are thus part of our selected datasets.

To cope with a limited number of unique training samples and imbalances in disease representation, we propose oversampling the underrepresented diseased donor samples. Unless carefully managed, oversampling can lead to overfitting or skewed class representations. However, in our model, as each training instance uses only a subset of available cells from a donor, oversampling from a single donor results in diverse training samples, mitigating traditional oversampling drawbacks and effectively increasing the variety of training instances. For example, with *n*_*genes*_ = 1000 and 2000 cells from a donor divided evenly across five cell types, there are over 10^590^ potential (unordered) cell combinations for selection. This strategy reduces the risk of overfitting to specific samples and helps to manage the issue of underrepresented diseases, although the distribution of cell types remains constant across oversampled instances. We introduce up to a 10 % randomized deviation in the stratification of cell types to vary this distribution slightly, while approximately still maintaining the original cell type distribution.

Furthermore, this randomized selection of cells for each new donor sample acts as a data augmentation strategy, mirroring the scRNA sequencing workflow where cells from a donor batch are randomly isolated and sequenced. Just as this randomness determines which cells are recorded in the datasets, our approach of randomly selecting 1023 cells per donor for each training epoch effectively simulates the natural variability of scRNA sequencing, enhancing the model’s robustness and representativeness.

### 3.3 Datasets and Preprocessing

The datasets utilized in our study are thoroughly detailed in Table 1, which lists each dataset along with its origin. This data has been quality-controlled, harmonized in terms of disease labels (e.g., grouping together “healthy”, “normal”, and “control”), and manually annotated to ensure standardization in terms of cell types and states, which are determined based on specific marker genes.

We preprocess the datasets used to train and test mcBERT via a three-step routine to facilitate efficient training. Beyond these three steps, no further preprocessing is necessary.

First, due to differences in sequencing methods across collections, not all genes are present in every dataset. However, mcBERT’s pipeline depends on the same set of gene’s across datasets and we opted to choose the 1000 most HVGs common to datasets of the same tissue type (see methods section). If the intersection of available HVGs across datasets contains fewer than 1000 genes, we greedily increase the number of HVGs considered from each dataset until the intersection reaches the desired threshold. This method avoids the need to impute genes that appear as HVGs in one dataset but are unsequenced in others, which could mislead mcBERT during training, potentially causing the model to learn imputation artifacts rather than true biological features.

Second, datasets comprising millions of cells can exceed the memory limits of the available RAM. To stay within memory limits, a single file for each donor is created, storing only the identified HVGs. This reduction not only conserves memory but also streamlines the data handling during the neural network training phase.

Third, each cell is normalized by the total counts over all remaining HVGs, and labels belonging to the same disease are harmonized, e.g., control, normal, and healthy are mapped to the “Healthy” class.

### 3.4 Inference

In the downstream analysis, the cosine similarities and distances between embeddings facilitate the comparison of cell donors. Additionally, for classifying a new patient with a potential disease, several methods are viable. Given that the diseases in the training dataset are labeled, these labels serve as anchor points during inference. A k-nearest neighbors (k-NN) algorithm can effectively classify a patient’s disease by identifying the most common disease within the patient’s local neighborhood of the training dataset. Moreover, to visualize the entire landscape of donor-based disease embeddings, a UMAP plot is created using the embedding cosine distances, providing a graphical representation of disease clustering and separation.

### 3.5 Evaluation Metrics

We evaluate the trained models on three different application tasks: (1) the direct patient similarity of the same and different disease class; (2) the disease classification using k-NN in relation to the training dataset; (3) the clustering using a hierarchical clustering approach.

The patient comparison task is the most natural task the model can be evaluated on as the contrastive loss used during fine-tuning trains the model specifically on this task. For numeric evaluation, we separately evaluate the mean cosine similarity of the test against the training samples for donors from the same class and from different classes, respectively. That is, we evaluate how close the trained model maps the donors’ embeddings next to other donors belonging to the same disease on the one hand and how far they are mapped away from different diseases on the other hand. The formal definitions of the two metrics are:

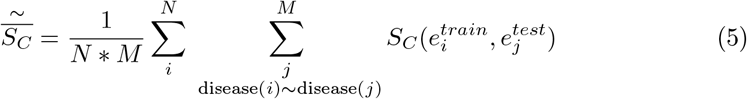

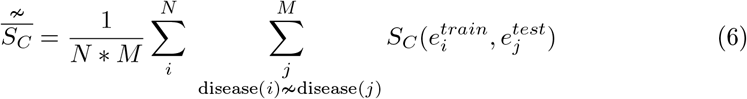

where *S*_*C*_ is the cosine similarity and *e* denotes the patient-level embedding. Ideally, a mean cosine similarity of 1 is achieved for patients with the same disease and 0 for different diseases.

For the second application task, the disease classification, the k-NN algorithm is used with k=5 and the cosine distance as the distance metric to determine the closest related donor. As k-NN relies on known data points to classify new samples, the samples from the training sets were taken and act as a database to classify the new donors. After classification by the k-NN algorithm, the accuracy can be determined. However, if the disease is not known in the training database, the accuracy is always 0 % and would distort the mcBERT metrics and is therefore not taken into account.

Lastly, to evaluate the global topology of our patient-level embedding, we employ a hierarchical clustering approach using agglomerative clustering with an average linkage criterion and cosine similarity as the distance metric and the number of unique diseases as the number of clusters. Similar to the patient-level distance evaluation of PILOT [20], the adjusted random index (ARI) [25] of the clustering is calculated to determine the clustering result. In combination to the hierarchical clustering, the silhouette coefficient [24] of the embedding is calculated using the cosine distance.

Evaluating the integration of single cells coming from different datasets is done as proposed by [27] using the local inverse Simpson’s index (LISI). The index LISI (iLISI) determines the integration regarding the dataset origin and is scaled between 0, worst integration, and 1, best integration. The cell-type LISI (cLISI) assesses the mixture of cell types and ideally maintains a value close to 1 before and after the integration of cells, which corresponds to well-separated cell-types in the embedding space.

## 4 Supplementary Material

### 4.1 Dataset Details

See Table 1.

### 4.2 mcBERT Architecture Details

The exact architectural details of mcBERT are shown in Table 2. Before processing the cells by the model, the pre-computed 1000 HVGs are selected. After the relatively lightweight dimensionality reduction step consisting of a single linear layer, a normalization layer, and a dropout layer with a dropout rate of 10 %, most of the computation falls to the transformer encoder. Here, each of the 12 consecutive transformer blocks contains 12 attention heads and has a hidden dimensionality of 288.

**Table 2:** The mcBERT pipeline is subdivided into the different embedding steps.

|  | Output size | Number of parameters | Details of Step |
| --- | --- | --- | --- |
| Pre-Selected Genes | (1024, 1002) | - | - |
| Cell Embedding | (1024, 288) | 288,864 | 1x Linear Layer |
|  | (1024, 288) | 576 | 1x LayerNorm |
|  | (1024, 288) | - | 1x Dropout, p=0.1 |
| Transformer Encoder | (1024, 288) | 25,282,944 | 12x Attention Blocks (12x Heads each) |
| (Pooling) | (1, 288) | - | 1x Global Average Pooling |
| $\Sigma$ | - | 25,572,384 | - |

The final global average pooling layer is used to reduce the contextualized cell matrix to a single patient-level vector. The learning of a donor’s cell correlations during the self-supervised training phase is not aimed at a single patient-level vector, that is why the pooling layer is omitted during the pre-training phase.

### 4.3 Metric Scores

Additionally to the detailed metrics of each cross-validation run outline in the supplemented Excel file, Table 3 contains the cross-validated metrics for the conducted experiments. For the first experiment shown in the table, only the reichart_2022 dataset was used. The leave-one-dataset-out cross-validation (LOOCV) averages over all runs where a different testing set was used and the model trained for the remaining datasets. The remaining experiments incorporate all selected datasets shown in Table 1 for the specified tissue.

**Table 3:**
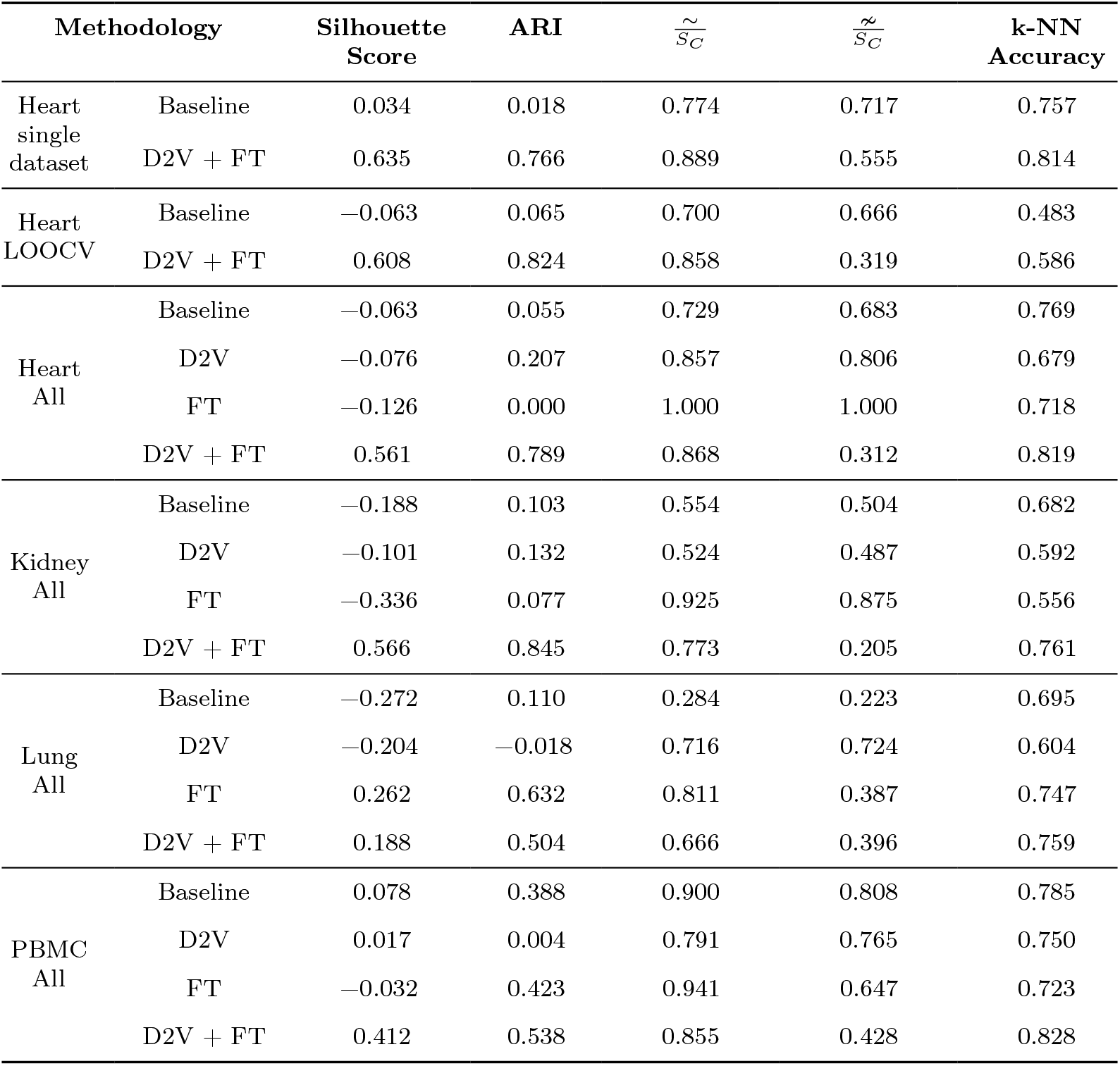
Averaged results of the cross-validation metrics of all experiments. D2V denotes the pre-training stage using Data2Vec, FT the fine-tuning and LOOCV the leave-one-dataset-out cross validation.

## Supplementary Information

All metrics of the conducted experiments are reported in an accompanied Excel file.

## Declarations

## Acknowledgments

S.H. is partly supported by the Leducq Early Career Investigator Award in the IMMUNO-FIB HF Network, CRU344, RWTH START, and Novo Nordisk STAR grants.

## Ethics Approval

Not applicable.

## Consent for Publication

Not applicable.

## Data Availability

The evaluation exclusively utilized publicly available datasets (see Table 1). All evaluation results are provided in a structured format in the supplementary Excel file (see “Supplementary Information”).

## Materials Availability

Not applicable.

## Code Availability

All software artifacts are available at https://github.com/COMSYS/mcBERT to support open science and to foster future research activities.

## Author Contributions

B.vQ. proposed the research approach, which was subsequently conceptualized and refined together with J.L., J.P., and S.H. Moreover, S.H. suggested the datasets to focus the evaluation on. T.B. contributed to human heart data integration. B.vQ. implemented mcBERT, conducted the evaluation, created the figures, and handled making data and code publicly available. B.vQ. drafted the manuscript, which was extensively revised by J.L., J.P., and S.H. Overall, J.L., J.P., and S.H. jointly supervised the work. All authors reviewed and approved the manuscript.

